# Weight loss-induced adipose macrophage memory improves local *Staphylococcus aureus* clearance in male mice

**DOI:** 10.1101/2024.08.03.606494

**Authors:** McArthur Bolden, Xenia D Davis, Edward R Sherwood, Julia K Bohannon, Heather L Caslin

**Affiliations:** Department of Health and Human Performance, University of Houston; Department of Anesthesiology, Vanderbilt University Medical Center; Department of Pathology, Microbiology, and Immunology, Vanderbilt University Medical Center

**Keywords:** Innate immune memory, trained innate immunity, weight loss, obesity, Staphylococcus aureus, infection

## Abstract

Different stimuli can induce innate immune memory to improve pathogen defense or worsen cardiometabolic disease. However, it is less clear if the same stimuli can induce both the protective and detrimental effects of innate immune memory. We previously showed that weight loss induces innate immune memory in adipose macrophages that correlates with worsened diabetes risk after weight regain. In this study, we investigated the effect of weight loss on macrophage cytokine production and overall survival in a mouse model of infection. Male C57Bl/6J mice were put on high-fat or low-fat diets over 18 weeks to induce weight gain or weight loss. Lean mice served as controls. All mice were then infected IV with 2.5×10^6 CFU *Staphylococcus aureus*. Tissues were collected from 10 mice/group at day 3 and the remaining animals were followed for survival. Weight gain mice had the highest blood neutrophils and the highest bacterial burden in the kidney. However, there was no significant difference in survival. The weight loss group had the highest plasma TNF-α and a significant reduction in bacterial burden in the adipose tissue that correlated with increased adipose macrophage cytokine production. Thus, weight loss-induced adipose macrophage memory may both improve local *S.aureus* clearance and worsen diabetes risk upon weight regain. Collectively, these findings support the notion that innate immune memory is an evolutionarily protective mechanism that also contributes to the development of cardiometabolic diseases.

## Introduction

Innate and adaptive immunity are critical components of host defense in the vertebrate immune system. Traditionally, the establishment of immune memory has been the fundamental role of lymphocytes in the adaptive immune system^1^. Following antigen-specific activation and memory formation, memory cells persist to mount a faster, more effective response if the same pathogen is encountered again. However, recent studies also show that the innate immune system can develop memory for a similar function^2,3^.

Innate immune memory, or “trained innate immunity”, develops in myeloid cells in response to an initial activation signal and enhances effector functions in response to a second, or subsequent, challenge^4^. Activation of innate receptors such TLR4 induce metabolic reprogramming and epigenetic imprinting to enhance effector functions like cytokine production and phagocytosis^4,5^. For example, the tuberculosis vaccine Bacille Calmette-Guérin (BCG), which was developed from *Mycobacterium bovis*, has been shown to have protective effects against not only *Mycobacterium tuberculosis*, but *Candida albicans*, *Schistosoma mansoni*, and malaria that can be attributed to innate immune memory^6–9^. Beta-glucan, a fungal polysaccharide, can also protect against *Pseudomonas aeruginosa, Staphylococcus aureus,* and *M. tuberculosis*^10–12^.

Additionally, the TLR4 agonist, monophosphoryl lipid A (MPLA), induces innate immune memory that protects against *S. aureus*, *P. aeruginosa,* and *Klebsiella pneumonia*^13–16^. Thus, innate immune memory is advantageous and distinct from that of antigen specific adaptive memory because of the range of stimuli that act as a primary signal and because secondary activation protects not only against reinfection, but against subsequent challenges with a different stimulus or pathogen.

Innate immune memory to BCG, beta-glucan, and MPLA ultimately boosts anti-microbial immunity. However, metabolic stimuli can also induce trained innate immunity that promotes chronic cardiometabolic disease. Oxidized-low density lipoprotein (ox-LDL) has been shown to induce innate immune memory in bone marrow derived macrophages^17,18^. Additionally, Western diet feeding worsens the induction and progression of atherosclerosis in mice^17,18^. Similarly, we have shown that palmitate or adipose tissue-conditioned media can induce innate immune memory in bone marrow-derived macrophages, with augmented cytokine production in response to lipopolysaccharide (LPS), lipoteichoic acid, poly(I:C), and beta-glucan^19^. In mice, weight loss increased maximal metabolism, LPS-induced cytokine production, and weight regain-induced cytokine production in adipose macrophages. Together, our data suggests that weight loss induces adipose macrophage memory that correlates with worsened glucose tolerance in our model of weight cycling.

The differences between the beneficial and detrimental outcomes of innate immune memory are still not well understood. In this study, we aimed to determine the effect of weight loss on murine survival and macrophage activation in a mouse model of infection.

## Materials and Methods

### Mice

Male C57BL/6J mice were purchased at 7 weeks of age from Jackson Labs (#000664). After 1 week of acclimation, all mice were placed on 9-week cycles of 60% high fat diet (Research Diets #D12492) or 10% low fat diet (Research Diets #D12450B) for a total of 18 weeks (see **Figure 1A**). Food and water were provided ad libitum throughout the study, and body weight and food intake were recorded weekly (see **Figure 1B&C**).

**Figure 1:**
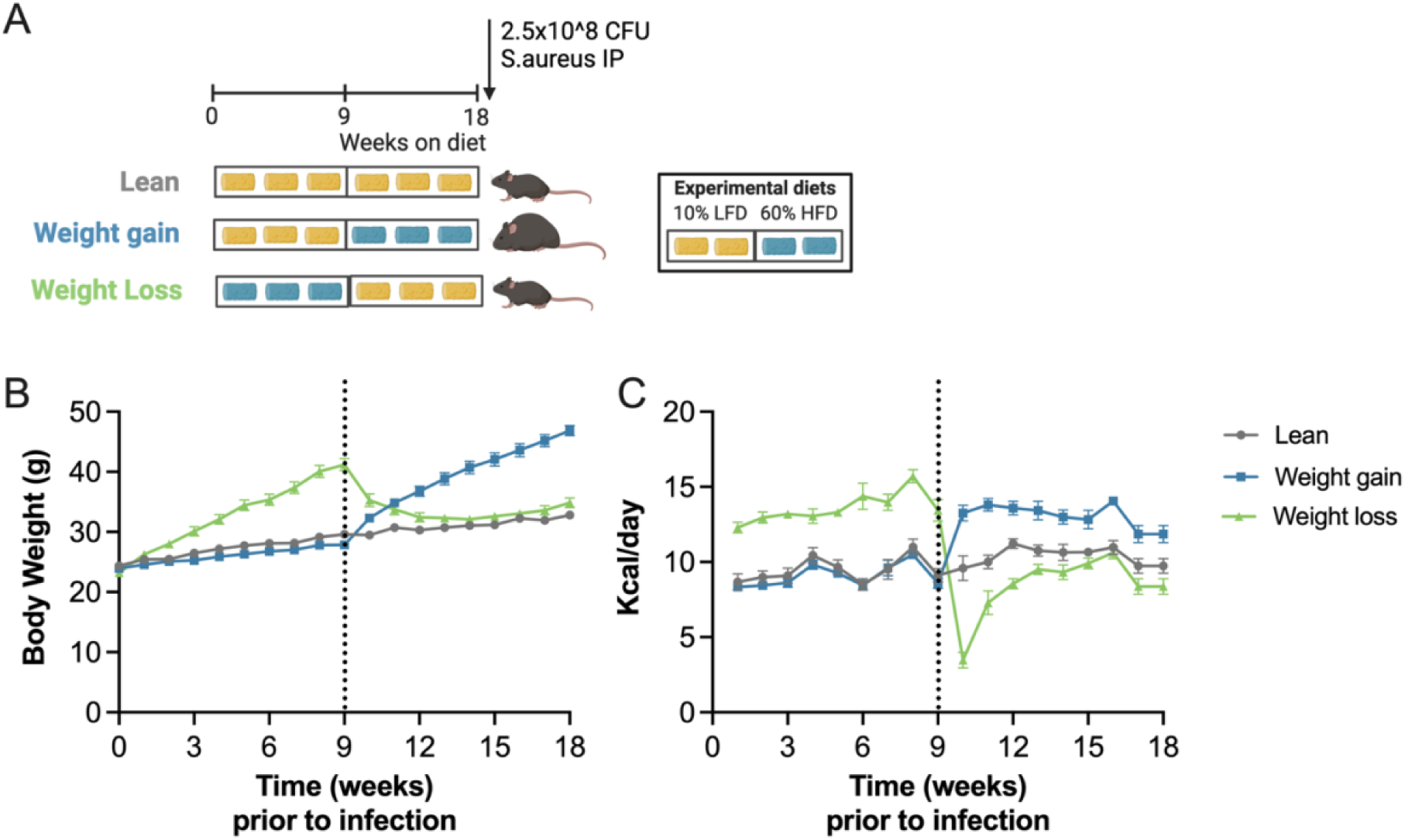
Schematic of diet-induced weight gain and loss prior to infection. A) Schematic of diet groups and infection created in Biorender. B) Body weight and C) food intake over time in lean, weight gain, and weight loss mice prior to infection. Error= SEM. n= 26-30/group.

All procedures were approved and performed at Vanderbilt University in compliance with the Vanderbilt University Institutional Animal Care and Use Committee. Vanderbilt University is accredited by the Association for Assessment and Accreditation of Laboratory Animal Care International.

### *Staphylococcus aureus* infection

Following 18 weeks on diet, *S. aureus*, obtained from American Type Culture Collection (ATCC #25923), was grown in tryptic soy broth for 22 hours at 37°C before centrifugation and resuspension in sterile saline. Mice were inoculated intravenously with 2.5 × 10^8 colony forming units (CFU); a dose chosen to elicit sufficient 3-day survival for all groups by a previous dose response curve. Body weight, rectal temperature, and survival were measured and recorded twice daily for the first week post-infection and at least once daily thereafter. Body condition was scored using a modified version the Murine Sepsis Score chart^20^ by the same individual each day. For our modification, the categories “level of consciousness” and “response to stimulus” were combined for one score and “respiration rate” and “respiration quality” were combined for one score. Ten mice/group were euthanized for tissue analysis at 3 days post-infection and 16-18 mice/group were followed for survival analysis. These mice were split between cohorts separated by about one year. Mice were euthanized if temperatures registered under 34°C or if the mice had apparent neurological symptoms (ie. a rolling response).

### CBC count and plasma analysis

At 3 days post-infection, 15 μL blood was collected in K3 EDTA Bio-One Mini Collect tubes (Greiner # 450532) and was sent to the Vanderbilt Translational Pathology Share Resource for complete blood counts (CBC). An additional 500 μL was suspended in heparin and plasma was frozen for cytokine analysis. TNF-α and IL-6 were measured by ELISA kits according to the manufacturers protocol (Biolegend #431304 & 430904 and RnD Systems #DY406).

### Tissue bacterial burden

At 3 days post-infection, a portion of the kidney, spleen, and gonadal fat pads were weighed and homogenized for bacterial burden. Samples were serially diluted in sterile saline and plated on tryptic soy agar overnight (37°C). After 24 hours of culture, colonies were counted, and total tissue burden/gram was calculated.

### Tissue fixation and histology

At 3 days post-infection, the kidney, spleen, and gonadal fat pads were harvested from 5 mice per group, fixed in 4% PFA, and paraffin embedded. The samples were sent to the Baylor College of Medicine Pathology and Histology Core, sectioned (∼8 μm), mounted, and two slides × 3 sections were stained with H&E. Two images were taken per animal from the upper right and lower left quadrants of the tissue on a Nikon Eclipse 80i upright microscope using the 4x objective. Representative samples are shown based on blinded image analysis and histological labeling was included as similarly described^21–24^.

### Adipose macrophage flow cytometry

Adipose cell isolation was completed as previously described^25^. Briefly, the gonadal adipose tissue remaining after a section was taken for histology was collected in PBS with 2% FBS. Tissue was minced and digested in 2 mg/mL type IV collagenase (Millipore Sigma, C4-BIOC) for 30 min at 37°C. Digested tissue was then vortexed and filtered through 100 μm filters. Adipocytes were discarded and stromal vascular fraction was lysed with ACK buffer and filtered again through a 35 μm filter for single cell suspensions.

For flow cytometry staining, Fc block was added to the isolated stromal vascular fraction for 10 min prior to fluorescent surface marker staining for 40 min at 4°C (antibodies and concentrations can be found in Table 1). Cells were then fixed and permeabilized for intracellular (cytoplasmic) proteins using the Foxp3/transcription factor staining buffer set (eBioscience /Invitrogen #00-5523-00). Cells were stained for intracellular proteins overnight at 4°C (see Table 1 for antibody information). Data was acquired on a MACSQuant10 (Miltenyi) and analyzed on FlowJo software (v10).

**Table 1.**

| **Antibody** | **Manufacturer** | **Catalog #** | **Dilution Factor** |
| --- | --- | --- | --- |
| FcBlock- Purified Rat Anti-Mouse CD16/CD32 | BD Pharmigen | 553141 | 200 |
| BV510 anti-mouse CD45 | Biolegend | 103137 | 400 |
| FITC anti-mouse CD11b | eBioscience | 11-0112-86 | 200 |
| PE-Cy7 anti-mouse CD9 | Biolegend | 124815 | 400 |
| APC-Cy7 anti-mouse F480 | Biolegend | 123118 | 200 |
| PE anti-mouse TNF-α | Biolegend | 506306 | 200 |
| APC anti-mouse IL-6 | Biolegend | 503810 | 200 |

### Statistical analyses

All data are presented as mean ± SEM and were analyzed using GraphPad Prism 10. Survival was analyzed using Kaplan-Meier curves with a Log-rank test to detect statistical significances between groups. Two-way ANOVA analyses were completed for 2+ groups across 2+ timepoints for body weight and food intake.

Bonferroni-corrected *post hoc* comparisons were used when appropriate to determine where differences occur between groups. One-way ANOVA analysis was performed for comparisons of multiple groups across one timepoint followed by Dunnett’s post hoc multiple comparison test. For correlation of plasma cytokines with temperature, a simple linear regression was conducted. A p-value of ≤ 0.05 was considered statistically significant.

## Results

### Weight gain and weight loss do not significantly affect infection survival

Weight loss worsens adipose inflammation during weight regain^19^, but the impact of weight loss prior to acute infection is much less clear. To determine the effect of weight loss on systemic *S.aureus* infection, mice were put on low fat or high fat diets as shown in **Figure 1A** then infected IV with 2.5 × 10^6 CFU *S.aureus*. Pre-infection, body weight, and food intake were measured weekly (see **Figure 1B&C**). Post-infection, body weight, temperature, and body condition score were recorded 1-2 times daily in all surviving mice as reported in **Figure 2A-C**. Moreover, survival was recorded in all mice (**Figure 2D**). There was no difference in survival between groups. The median survival times were 2.25, 2.75, and 3.75 for the lean, weight gain, and weight loss groups, respectively.

**Figure 2:**
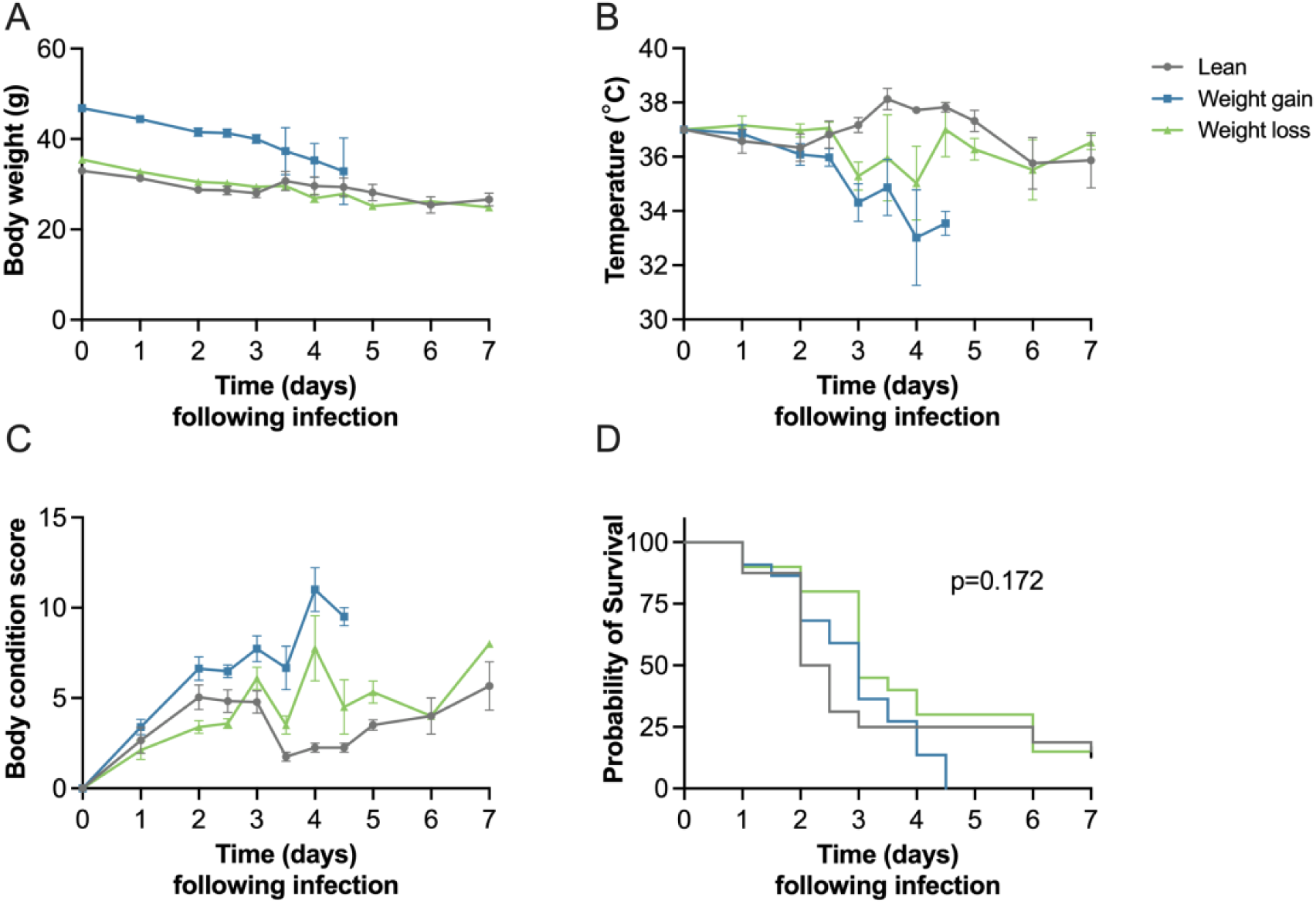
Weight gain worsens body condition score, but survival is unchanged between groups. A) Body weight, B) temperature, and C) body condition score following infection in lean, weight gain, and weight loss following infection. D) Survival over one week. Error= SEM. n=21-24/ group. Statistics for survival by Log-rank analysis.

Weight gain increases neutrophils in the blood and weight loss increases plasma TNF-α Survival in sepsis is dependent upon having sufficient immune activation to clear the pathogen but restraining inflammation enough to protect the host. To determine the effect of weight loss on white blood cells, complete blood counts were analyzed from 10 mice/group at day 3 after infection as reported in **Figure 3A**. When compared to the lean group, the only observed differences were an increase in neutrophil number and a corresponding increase in total white blood cells in the weight gain group. Additionally, plasma cytokines were analyzed by ELISA. There was a significant increase in TNF-α in the weight loss group compared with the lean group (**Figure 3B**), but no difference in IL-6 between groups (**Figure 3C**). Interestingly, there was a negative correlation between IL-6 and temperature on day 3 (**Figure 3C**), which was not observed with TNF-α (**Figure 3B**).

**Figure 3:**
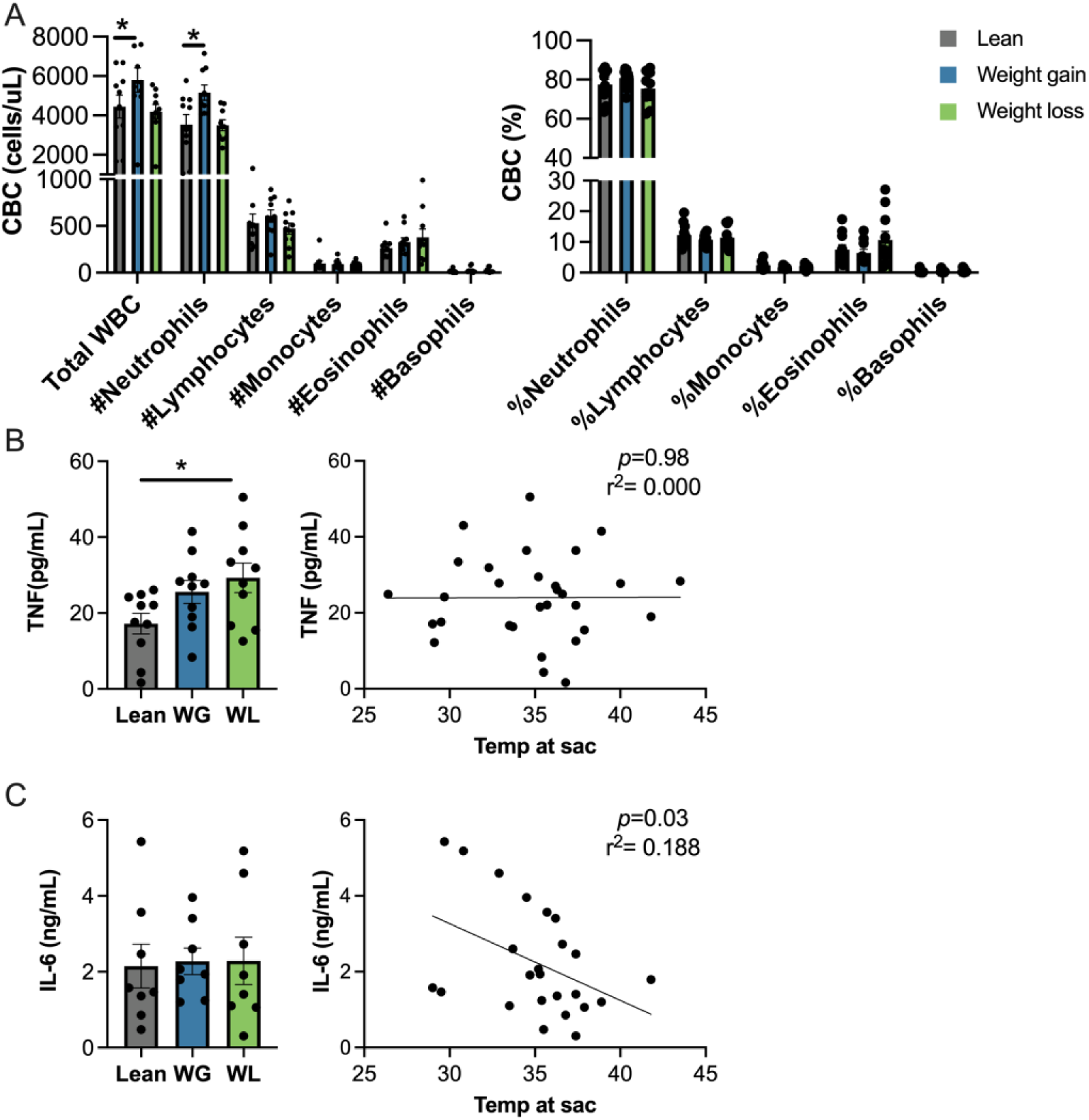
Weight gain increases plasma neutrophils while weight loss increases plasma. TNF-α. Complete blood counts by A) cells/ uL or B) percentage in plasma from lean, weight gain, and weight loss mice at 3 days following infection. C) TNF-α and D) IL-6 in the plasma as measured in each group by ELISA at 3 days following infection and the association between plasma cytokine concentrations and temperature at day 3 were computed. Error= SEM. n=8-10/group. Statistical analysis by one-way ANOVA and Dunnett’s post hoc analyses (A-D) and simple linear regression (C-D). * p<0.05.

### Weight loss improves bacterial clearance in the adipose tissue which corresponds to worsened increased macrophage activation

Local bacterial clearance can affect organ function and ultimately survival. To determine the impact of weight loss on bacterial clearance in the tissue, we measured bacterial burden in kidney, spleen, and adipose tissue at day 3. Weight gain increased bacterial burden in the kidney compared with the lean and weight loss groups (**Figure 4A**). There were abscesses of immune infiltration and bacterial colonies in the kidney sections from the weight gain animals. Hyperemic interstitial capillaries and interstitial hemorrhage were seen in all groups (representative images shown in **Figure 4B**). In the spleen, bacterial burden was not significantly different between groups (**Figure 5A**). An expansion of white pulp area was observed in all groups (representative images shown in **Figure 5B**). Notably, there was a significant reduction in bacterial burden in the adipose tissue from weight loss mice compared with weight gain mice (**Figure 6A**). Additionally, there was a trend towards improved bacterial clearance in the weight loss compared with lean mice. All groups showed hyperemic capillaries, and the weight loss group had the largest density of crown-like structures (representative images shown in **Figure 6B**).

**Figure 4:**
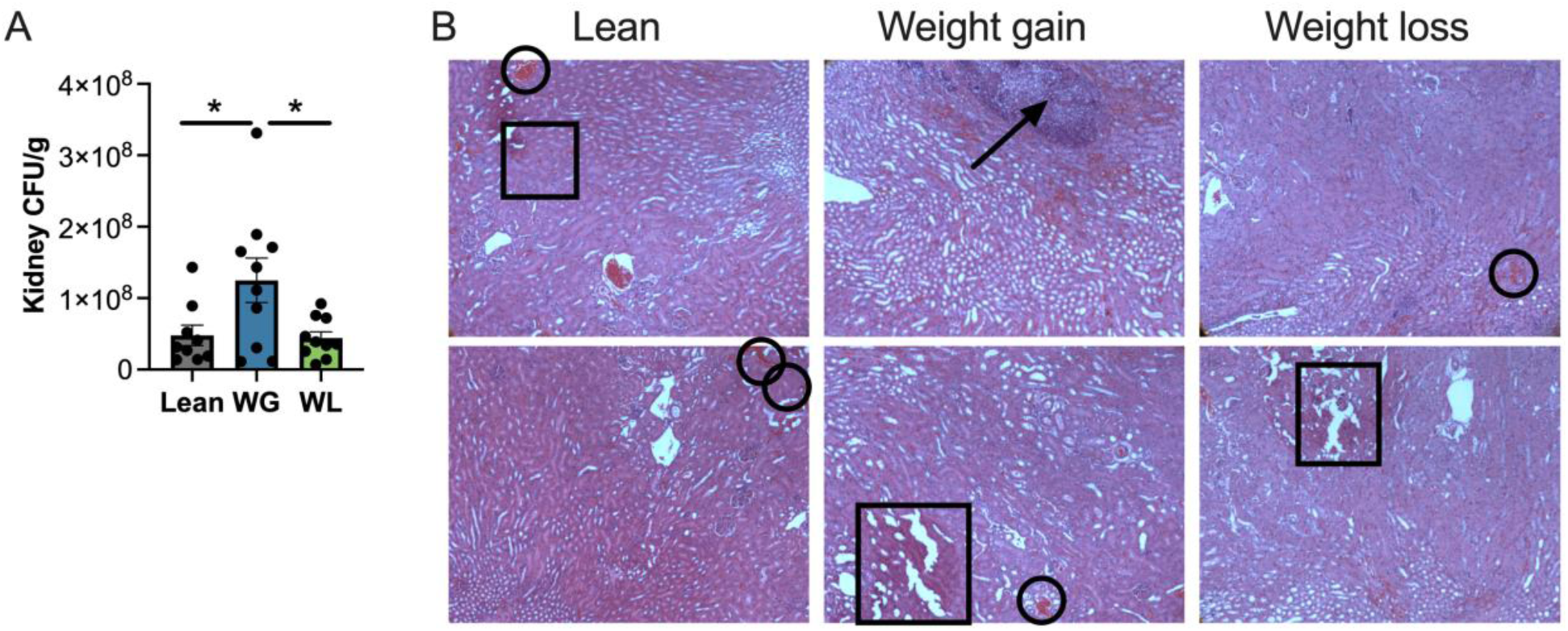
Weight gain worsens bacterial clearance in the kidney. A) Bacterial clearance by colony forming units/ gram tissue in kidney from lean, weight gain, and weight loss mice 3 days following infection. B) Two representative images from each group. The arrow(s) signify immune infiltrate and a bacterial colony abscess, the circle(s) signify hyperemic interstitial capillaries, and the square(s) signify interstitial hemorrhage and tissue breakdown. Error= SEM. n=9-10/group. Statistical analysis by one-way ANOVA and Dunnett’s post hoc analyses. * p<0.05.

**Figure 5:**
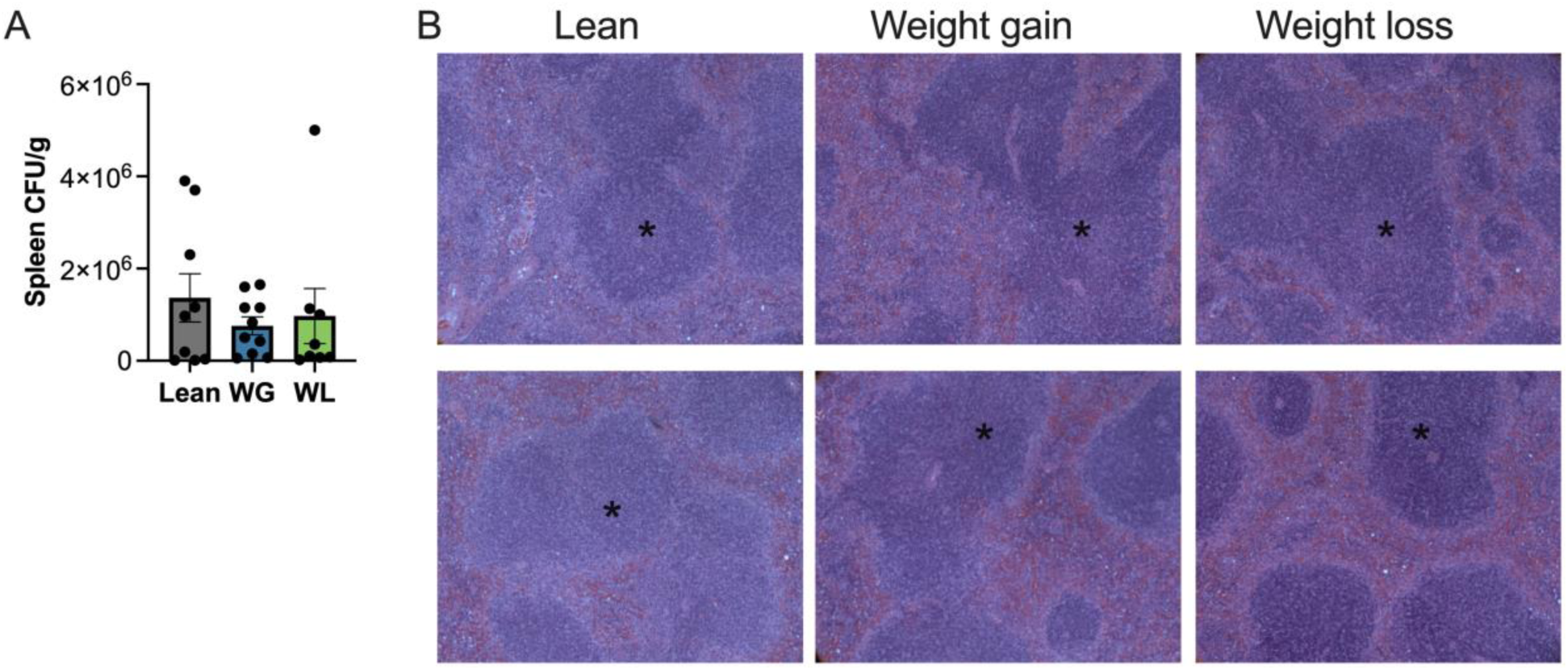
Weight change has no impact on bacterial clearance in the spleen. A) Bacterial clearance by colony forming units/ gram tissue in spleen from lean, weight gain, and weight loss mice 3 days following infection. B) Two representative images from each group. The arrow(s) signify a region of white pulp. Error= SEM. n=8-10/group. Statistical analysis by one-way ANOVA and Dunnett’s post hoc analyses.

**Figure 6:**
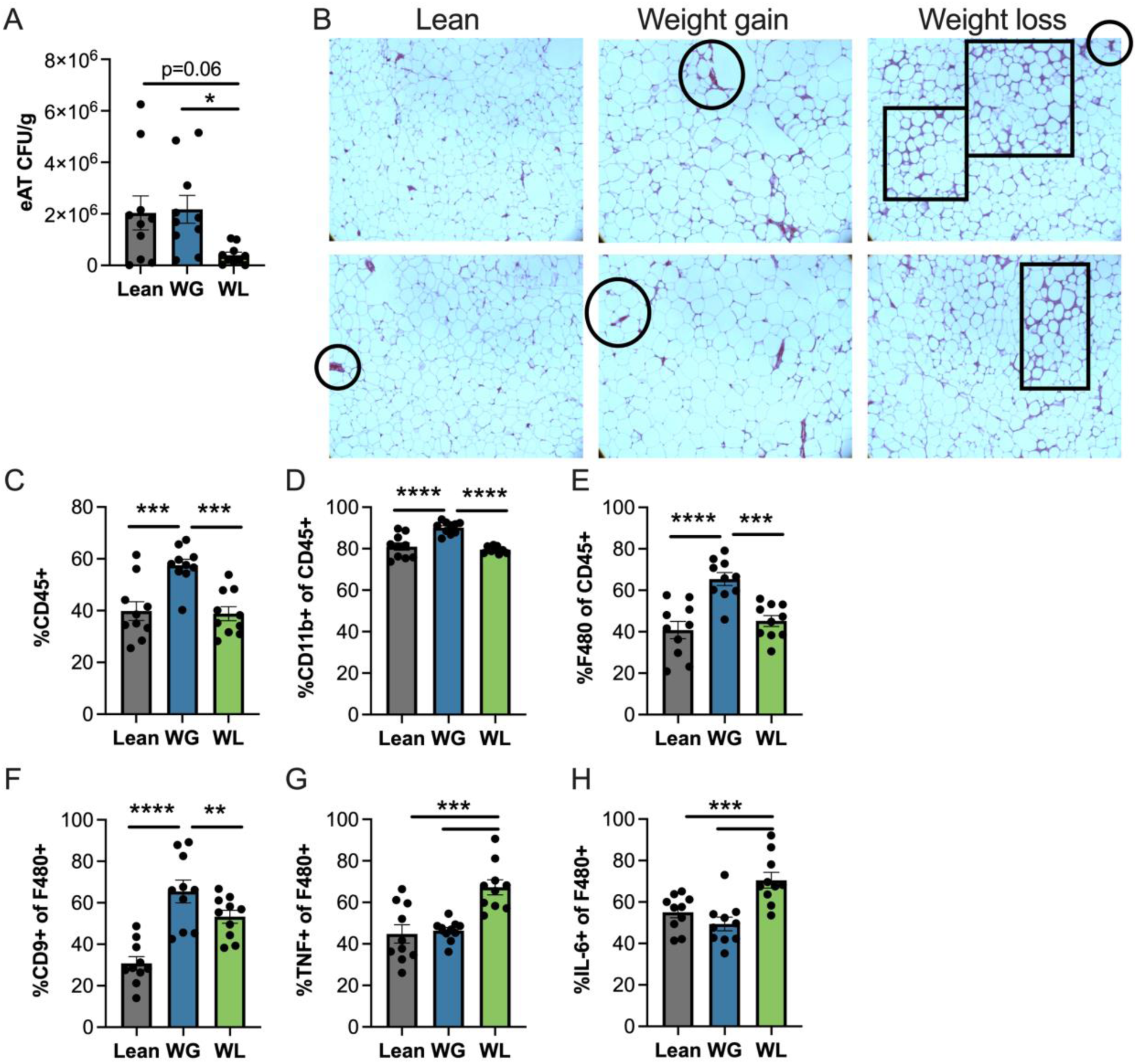
Weight loss improves bacterial clearance in the adipose, which corresponds with heightened macrophage cytokine production. A) Bacterial clearance by colony forming units/ gram tissue in adipose from lean, weight gain, and weight loss mice 3 days following infection. B) Two representative images from each group. The circle(s) signify hyperemic interstitial capillaries, and the square(s) signify crown-like structures. C) %CD45+, D) %CD11b+, E) %F4/80+, and F) %CD9+ populations by flow cytometry from each group. G) % TNF-α + and H) %IL-6+ F4/80+ macrophages by intracellular flow cytometry. Error= SEM. n=9-10/group. Statistical analysis by one-way ANOVA and Dunnett’s post hoc analyses. * p<0.05, **p<0.01, ***p<0.001, ****p<0.0001

One function of the innate immune system in infection involves the production of inflammatory cytokines. Cytokines like TNF-α and IL-6 can induce immune cell recruitment and activation and can induce iron sequestration to help control infection by pathogens like *S.aureus*^26,27^. Because weight loss mice had improved adipose tissue bacterial clearance and elevated plasma TNF-α, we were interested to understand how weight loss impacts the proportion of adipose macrophages and adipose macrophage cytokine production to *S.aureus* following weight loss. Weight gain increased the proportion of immune cells, myeloid cells, macrophages, and CD9+ macrophages (as a proxy for lipid-associated macrophages^28^; **Figure 6C-F**). However, these results are likely a product of the weight gain itself, and not the infection, as similar findings are often reported in adipose tissue of obese, non-infected mice^19,28,29^. Interestingly, the percent of TNF-α + and IL-6+ adipose macrophages was highest in the weight loss group, corresponding with the improved bacterial clearance.

## Discussion

Stimuli like BCG, beta-glucan, and MPLA can induce innate immune memory which improves pathogen defense^4,30^. Ox-LDL, palmitate, and hyperglycemia also induce innate immune memory, however, this worsens the development of cardiometabolic disease^17,19,31^. How individual stimuli impact both pathogen defense and chronic inflammatory disease is less clear. We previously demonstrated that weight loss can induce adipose macrophage memory, defined by heightened metabolism and heightened cytokine production to LPS and weight regain^19^. This inflammatory state correlated with worsened diabetes risk upon weight regain. In this study, we sought to understand the impact of weight loss-induced innate immune memory on the response to *S.aureus* infection. We found that weight loss increases bacterial clearance and adipose macrophage cytokine production specifically in the adipose tissue. Thus, while weight loss-induced adipose macrophage memory may worsen diabetes risk following weight regain, the same stimuli improves *S.aureus* clearance and pathogen defense in the local adipose tissue.

At the systemic level, we found that diet-induced weight gain and weight loss had no impact on survival to *S.aureus.* However, it should be noted that between 1-3 days, the weight gain and weight loss animals seemed to fare better, but by day 4-5, the weight gain animals had much worse sepsis scores and seemingly worse survival.

Thus, there is an interesting relationship between weight and response to infection that is also supported in the literature. In humans, some report an obesity paradox, in which sepsis patients who meet the criteria for overweight and obese have lower mortality than lean individuals^32–34^. This may be due to patients with higher adiposity having higher metabolic reserves, higher secretion of anti-inflammatory mediators by adipose tissue, and higher lipoprotein concentrations to bind to bacterial LPS to dampen the early cytokine storm^35,36^. However, not all studies report better outcomes with higher adiposity, especially in older or more severe sepsis patients^34,37,38^. In animal models, differential outcomes may be due to differences in total time on high fat diet, specific model, and specific outcome^39–43^. Thus, there is still much to learn about the impact of adipose tissue on infection responses. Because we observed substantial within-group variability and potential between-group differences in survival at early vs. later timepoints, future studies may benefit from a larger sample size and analysis of tissues at both an earlier and a later timepoint.

Notably, the biggest effect observed in our study was a local improvement in bacterial clearance in the adipose tissue which corresponded to greater macrophage cytokine production following weight loss. Interestingly, a saturated fatty acid-rich diet and systemic palmitate administration also augment systemic inflammation to LPS and enhance clearance of *Candida albicans*^44^. Thus, we postulate that our effect is driven by weight loss-induced lipolysis. Additionally, we postulate that improved bacterial clearance is due to the enhanced macrophage cytokine production observed in our model. Interestingly, CD8+ memory T cells exhibit increased exhaustion, impaired metabolism, and impaired function following weight gain which is not reversed, and actually worsens, with weight loss^29,45^. Thus, it’s unlikely that the improved bacterial clearance is due to CD8+ memory T cells in our studies. We also cannot rule out the potential impact of weight loss on other macrophage functions like phagocytosis and ROS production or other immune (and non-immune^46^) populations. However, it’s important to note that improved bacterial clearance was localized to the adipose tissue. Our previous data showed that weight loss-induced innate immune memory was localized to adipose macrophages, but not liver or peritoneal macrophages^19^. Thus, it’s plausible that enhanced macrophage inflammation and improved bacterial clearance in weight loss mice being localized to the adipose suggest a causal relationship.

On another note, not all inducers of innate immune memory increase cytokine production. While we see increased TNF-α in adipose macrophages and the plasma with weight loss-induced memory, MPLA does not increase inflammatory cytokine production by macrophages and may improve pathogen defense via other functions like phagocytosis, ROS production, and chemokine production^10,13^. Thus, while there are many similarities between different innate immune memory stimuli, there may be specific effects related to adiposity due to the presence of chronic low-grade inflammation, lipids, or other hormonal signals. Future work should further unravel the impact of weight loss on other macrophage functions like phagocytosis, ROS production, and chemokine production. Interestingly, our previous study also showed that adipose macrophages in multiple fat depots and liver macrophages had increased basal cytokine production following weight regain^19^, suggesting systemic induction of innate immune memory following weight regain. It would be particularly interesting to look at the effect of weight regain on local and systemic bacterial clearance, which may also provide insight into human studies.

Taken together, these studies suggest that weight loss-induces innate immune memory in adipose tissue macrophages that correlates with not only worsened diabetes risk, but improved *S.aureus* clearance. Thus, while different innate immune memory stimuli improve pathogen defense, but worsens chronic disease, it seems that individual stimuli may specifically contribute to both the beneficial and detrimental outcomes as well. This knowledge may shed light on how an evolutionarily protective innate immune response may contribute to the development of cardiometabolic diseases in our modern environment.

## Acknowledgements

We’d like to acknowledge Dr. Leigh Leasure at the University of Houston for the use of her microscope for histology image analysis, as well as the Vanderbilt Mouse Metabolic Phenotypic Center (supported by DK135073), the Vanderbilt Translational Pathology Share Resource Core (supported by 2P30 CA068485 and 5U24DK059637), the Baylor College of Medicine Pathology and Histology Core (supported by NCI-CA125123).

## Notes

### Competing Interest Statement

The authors have declared no competing interest.

## References

1. Bonilla FA, Oettgen HC. Adaptive immunity. Journal of Allergy and Clinical Immunology. 2010;125(2):S33–S40. doi:10.1016/j.jaci.2009.09.017

2. Tercan H, Riksen NP, Joosten LAB, Netea MG, Bekkering S. Trained Immunity: Long-Term Adaptation in Innate Immune Responses. Arterioscler Thromb Vasc Biol. 2021;41(1):55–61. doi:10.1161/ATVBAHA.120.314212

3. Netea MG, Domínguez-Andrés J, Barreiro LB, et al. Defining trained immunity and its role in health and disease. Nat Rev Immunol. 2020;20(6):375–388. doi:10.1038/s41577-020-0285-6

4. Netea MG, Joosten LAB, Latz E, et al. Trained immunity: A program of innate immune memory in health and disease. Science. 2016;352(6284):aaf1098–aaf1098. doi:10.1126/science.aaf1098

5. Ochando J, Mulder WJM, Madsen JC, Netea MG, Duivenvoorden R. Trained immunity — basic concepts and contributions to immunopathology. Nat Rev Nephrol. 2023;19(1):23–37. doi:10.1038/s41581-022-00633-5

6. Chen J, Gao L, Wu X, et al. BCG-induced trained immunity: history, mechanisms and potential applications. J Transl Med. 2023;21:106. doi:10.1186/s12967-023-03944-8

7. Tribouley J, Tribouley-Duret J, Appriou M. [Effect of Bacillus Callmette Guerin (BCG) on the receptivity of nude mice to Schistosoma mansoni]. C R Seances Soc Biol Fil. 1978;172(5):902–904.

8. Van’t Wout JW, Poell R, Van Furth R. The Role of BCG/PPD-Activated Macrophages in Resistance against Systemic Candidiasis in Mice. Scandinavian Journal of Immunology. 1992;36(5):713–720. doi:10.1111/j.1365-3083.1992.tb03132.x

9. Walk J, de Bree LCJ, Graumans W, et al. Outcomes of controlled human malaria infection after BCG vaccination. Nat Commun. 2019;10(1):874. doi:10.1038/s41467-019-08659-3

10. Stothers CL, Burelbach KR, Owen AM, et al. Beta-glucan Induces Distinct and Protective Innate Immune Memory in Differentiated Macrophages. J Immunol. 2021;207(11):2785–2798. doi:10.4049/jimmunol.2100107

11. Moorlag SJCFM, Khan N, Novakovic B, et al. β-Glucan Induces Protective Trained Immunity against Mycobacterium tuberculosis Infection: A Key Role for IL-1. Cell Rep. 2020;31(7):107634. doi:10.1016/j.celrep.2020.107634

12. Di Luzio NR, Williams DL. Protective effect of glucan against systemic Staphylococcus aureus septicemia in normal and leukemic mice. Infect Immun. 1978;20(3):804–810.

13. Fensterheim BA, Young JD, Luan L, et al. The TLR4 Agonist Monophosphoryl Lipid A Drives Broad Resistance to Infection via Dynamic Reprogramming of Macrophage Metabolism. The Journal of Immunology. 2018;200(11):3777–3789. doi:10.4049/jimmunol.1800085

14. Romero CD, Varma TK, Hobbs JB, Reyes A, Driver B, Sherwood ER. The Toll-Like Receptor 4 Agonist Monophosphoryl Lipid A Augments Innate Host Resistance to Systemic Bacterial Infection. Infection and Immunity. 2011;79(9):3576. doi:10.1128/IAI.00022-11

15. Hernandez A, Zhou J, Bohannon JK, et al. INTRAPULMONARY TREATMENT WITH A NOVEL TLR4 AGONIST CONFERS PROTECTION AGAINST KLEBSIELLA PNEUMONIA. Shock. 2022;58(4):295–303. doi:10.1097/SHK.0000000000001977

16. Owen AM, Fults JB, Patil NK, Hernandez A, Bohannon JK. TLR Agonists as Mediators of Trained Immunity: Mechanistic Insight and Immunotherapeutic Potential to Combat Infection. Frontiers in Immunology. 2020;11. doi:10.3389/fimmu.2020.622614

17. Bekkering S, Quintin J, Joosten LAB, van der Meer JWM, Netea MG, Riksen NP. Oxidized low-density lipoprotein induces long-term proinflammatory cytokine production and foam cell formation via epigenetic reprogramming of monocytes. Arterioscler Thromb Vasc Biol. 2014;34(8):1731–1738. doi:10.1161/ATVBAHA.114.303887

18. Keating ST, Groh L, Thiem K, et al. Rewiring of glucose metabolism defines trained immunity induced by oxidized low-density lipoprotein. J Mol Med (Berl). 2020;98(6):819–831. doi:10.1007/s00109-020-01915-w

19. Caslin HL, Cottam MA, Piñon JM, Boney LY, Hasty AH. Weight cycling induces innate immune memory in adipose tissue macrophages. Front Immunol. 2023;13:984859. doi:10.3389/fimmu.2022.984859

20. Shrum B, Anantha RV, Xu SX, et al. A robust scoring system to evaluate sepsis severity in an animal model. BMC research notes. 2014;7(1):233.

21. Silva-Santana G, Lenzi-Almeida K, Fern, et al. Mice Infection by Methicillin-Resistant Staphylococcus Aureus from Different Colonization Sites in Humans Resulting in Difusion to Multiple Organs. Journal of Clinical & Experimental Pathology. 2016;6(3):1.

22. Balgradean M, Ceausu M, Cinteza E, et al. Death caused by hemolytic-uremic syndrome. Case series. RJLM. 2013;21(3):185–192. doi:10.4323/rjlm.2013.185

23. Eren Sadioglu R, Eyupoglu S, Kiremitci S, Birengel S, Keven K. Two patients, two viruses and multiple sites of injury in the kidney. J Nephrol. 2021;34(1):263–265. doi:10.1007/s40620-020-00838-6

24. Rauch S, DeDent AC, Kim HK, Bubeck Wardenburg J, Missiakas DM, Schneewind O. Abscess Formation and Alpha-Hemolysin Induced Toxicity in a Mouse Model of Staphylococcus aureus Peritoneal Infection. Weiser JN, ed. Infect Immun. 2012;80(10):3721–3732. doi:10.1128/IAI.00442-12

25. Orr JS, Kennedy AJ, Hasty AH. Isolation of Adipose Tissue Immune Cells. JoVE (Journal of Visualized Experiments). 2013;(75):e50707. doi:10.3791/50707

26. Youn C, Pontaza C, Wang Y, et al. Neutrophil-intrinsic TNF receptor signaling orchestrates host defense against Staphylococcus aureus. Science Advances. 2023;9(24):eadf8748. doi:10.1126/sciadv.adf8748

27. Pidwill GR, Gibson JF, Cole J, Renshaw SA, Foster SJ. The Role of Macrophages in Staphylococcus aureus Infection. Front Immunol. 2021;11. doi:10.3389/fimmu.2020.620339

28. Jaitin DA, Adlung L, Thaiss CA, et al. Lipid-Associated Macrophages Control Metabolic Homeostasis in a Trem2-Dependent Manner. Cell. 2019;178(3):686–698.e14. doi:10.1016/j.cell.2019.05.054

29. Cottam M, Caslin H, Winn N, Hasty A. Multiomics reveals persistence of obesity-associated immune cell phenotypes in adipose tissue during weight loss and subsequent weight regain. Nature Communications. 2022;13(1):2950. doi:10.1038/s41467-022-30646-4

30. Sherwood ER, Burelbach KR, McBride MA, et al. Innate Immune Memory and the Host Response to Infection. J Immunol. 2022;208(4):785–792. doi:10.4049/jimmunol.2101058

31. Robinson KA, Akbar N, Baidžajevas K, Choudhury RP. Trained immunity in diabetes and hyperlipidemia: Emerging opportunities to target cardiovascular complications and design new therapies. FASEB J. 2023;37(11):e23231. doi:10.1096/fj.202301078R

32. Elkan M, Kofman N, Minha S, Rappoport N, Zaidenstein R, Koren R. Does the “Obesity Paradox” Have an Expiration Date? A Retrospective Cohort Study. J Clin Med. 2023;12(21):6765. doi:10.3390/jcm12216765

33. Yeo HJ, Kim TH, Jang JH, et al. Obesity Paradox and Functional Outcomes in Sepsis: A Multicenter Prospective Study. Crit Care Med. 2023;51(6):742–752. doi:10.1097/CCM.0000000000005801

34. Dramé M, Godaert L. The Obesity Paradox and Mortality in Older Adults: A Systematic Review. Nutrients. 2023;15(7):1780. doi:10.3390/nu15071780

35. Antonopoulos AS, Tousoulis D. The molecular mechanisms of obesity paradox. Cardiovascular Research. 2017;113(9):1074–1086. doi:10.1093/cvr/cvx106

36. Donini LM, Pinto A, Giusti AM, Lenzi A, Poggiogalle E. Obesity or BMI Paradox? Beneath the Tip of the Iceberg. Front Nutr. 2020;7:53. doi:10.3389/fnut.2020.00053

37. Kalani C, Venigalla T, Bailey J, Udeani G, Surani S. Sepsis Patients in Critical Care Units with Obesity: Is Obesity Protective? Cureus. 2020;12(2). doi:10.7759/cureus.6929

38. Ng PY, Eikermann M. The obesity conundrum in sepsis. BMC Anesthesiol. 2017;17(1):147, s12871-017-0434-z. doi:10.1186/s12871-017-0434-z

39. Kaplan JM, Nowell M, Lahni P, O’Connor M, Hake PW, Zingarelli B. Short-Term High Fat Feeding Increases Organ Injury and Mortality After Polymicrobial Sepsis. Obesity (Silver Spring, Md). 2012;20(10):1995. doi:10.1038/oby.2012.40

40. Kaplan JM, Nowell M, Lahni P, Shen H, Shanmukhappa SK, Zingarelli B. Obesity enhances sepsis-induced liver inflammation and injury in mice. Obesity (Silver Spring). 2016;24(7):1480–1488. doi:10.1002/oby.21504

41. Siegl D, Annecke T, Johnson BL III, et al. Obesity-induced Hyperleptinemia Improves Survival and Immune Response in a Murine Model of Sepsis. Anesthesiology. 2014;121(1):98–114. doi:10.1097/ALN.0000000000000192

42. Shapiro NI, Khankin EV, Van Meurs M, et al. Leptin exacerbates sepsis-mediated morbidity and mortality. J Immunol. 2010;185(1):517–524. doi:10.4049/jimmunol.0903975

43. Eng M, Suthaaharan K, Newton L, Sheikh F, Fox-Robichaud A. Sepsis and obesity: a scoping review of diet-induced obesity murine models. Intensive Care Med Exp. 2024;12:15. doi:10.1186/s40635-024-00603-0

44. Seufert AL, Hickman JW, Traxler SK, et al. Enriched dietary saturated fatty acids induce trained immunity via ceramide production that enhances severity of endotoxemia and clearance of infection. Elife. 2022;11:e76744. doi:10.7554/eLife.76744

45. Rebeles J, Green WD, Alwarawrah Y, et al. Obesity-Induced Changes in T-Cell Metabolism Are Associated With Impaired Memory T-Cell Response to Influenza and Are Not Reversed With Weight Loss. J Infect Dis. 2019;219(10):1652–1661. doi:10.1093/infdis/jiy700

46. Caputa G, Matsushita M, Sanin DE, et al. Intracellular infection and immune system cues rewire adipocytes to acquire immune function. Cell Metabolism. 2022;34(5):747–760.e6. doi:10.1016/j.cmet.2022.04.008

